# The interplay between *Caenorhabditis elegans* larval development and Orsay virus infection

**DOI:** 10.1101/2025.05.23.655711

**Authors:** Izan Melero, Victoria G. Castiglioni, María J. Olmo-Uceda, María Olmedo, Rubén González, Santiago F. Elena

## Abstract

*Caenorhabditis elegans* development can be altered by its interactions with pathogens, such as bacteria^1,2^ or fungi^3^. However, the impact of viral infections on the nematode’s development remains largely unexplored, and conversely, whether molting periods affect viral replication is unknown. Studying infections with Orsay virus (OrV), the only known natural *C. elegans* virus^4^, we investigated the role of molting on virus replication dynamics and the impact of OrV infection on *C. elegans* larval development. We found that OrV replicates during *C. elegans* molting periods. Indeed, nematodes inoculated near molt initiation exhibited higher OrV viral loads and the virus infected more intestinal cells. Moreover, nematodes inoculated right after hatching accelerated the start of the first molt and lengthened the second larval stage, effectively resynchronizing normal developmental timing. Overall, our work enhances understanding of OrV dynamics in *C. elegans* and elucidates the interplay between viral infection and nematode development.

**HIGHLIGHTS:**

- rsay virus replication peaks during molting periods, with larvae acquiring virus later in the first larval stage producing higher viral loads and more widespread intestinal infection than larvae acquiring it right after hatching.
- infection alters the expression of key developmental regulators, particularly those with oscillatory profiles synchronized with molting cycles (such as *let-7*, *lin-42* and *nhr-23*).
- nematodes display temporal plasticity in development: accelerating the first molt while extending the second larval stage, ultimately resynchronizing their developmental timeline.
- is a bidirectional interaction between Orsay virus infection and *C. elegans* development: molting appears to transiently weaken antiviral defenses, while host development adapts to infection through temporal plasticity.

## RESULTS

### OrV replicates during molting, and molt initiation does not drive viral load decay

*C. elegans* develops through four larval stages (L1 to L4) before reaching adulthood^5^ (Figure S1). Between each stage, the nematode undergoes molting, a lethargic phase where animals renew their cuticle while stopping feeding and entering into a sleep-like state that limits locomotion^6,7^. A previous study on OrV dynamics throughout *C. elegans* larval development reported a viral load decay after reaching a maximum ∼12 hours post-inoculation (hpi)^8^. To investigate whether this viral decline results from the onset of molting, which starts around 15 hours post-hatching (hph) at 20 °C, we inoculated L1 nematodes immediately after hatching (0 hph) or at 6 hph. Samples were collected every 2 h for 24 h, covering the first molting period, and OrV viral load was measured by RT-qPCR to track replication dynamics. The inoculation at 6 hph was chosen to allow sufficient time for exponential viral replication before molt initiation (Figure S1), as this is the minimum time OrV requires to start exponentially replicating in intestinal cells under our conditions (see Figure 1B in Castiglioni et al.^8^).

**Figure 1.**
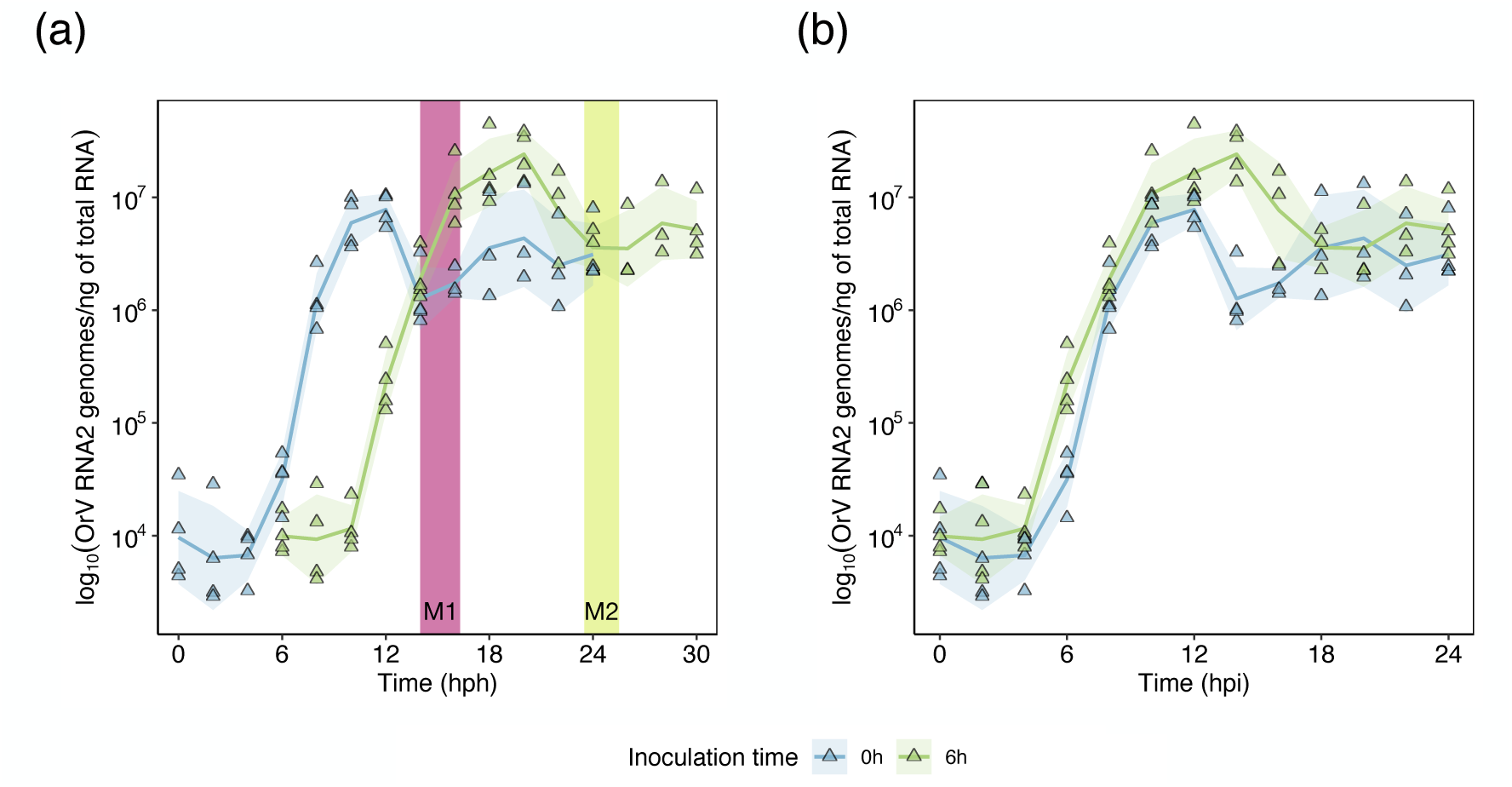
OrV accumulation dynamics. **(a)** Log_10_ RNA2 genomes/ng of total RNA measured every 2 h during 24 hpi. The abscissa shows hph. The first molting period (M1) is highlighted by a pink rectangle, and the second molting period (M2) by a lime green one. Blue lines and symbols represent the viral load for nematodes inoculated at 0 hph while green ones represent those inoculated at 6 hph. Shadowed areas represent ±1 standard deviation (SD). **(b)** Same data as in (a) but rescaled according to hpi.

We observed that the viral load of animals inoculated at 0 hph peaked around 12 hpi and subsequently declined, replicating previous observations^8^. In contrast, for animals inoculated at 6 hph, the molting period aligned with the phase of exponential viral replication (Figure 1A). This demonstrates that OrV can replicate within intestinal cells even while *C. elegans* undergoes molting. Furthermore, molt initiation not necessarily led to a viral load decline, and inoculating nematodes closer to the molting onset (6 hph) did not accelerate any decrease in viral titer. This suggests that the observed viral load decay after ∼12 hpi is likely results from factors other than the onset of molting.

### Inoculation of late L1 animals results in a higher and more prolonged first wave of OrV replication compared with inoculation of freshly-hatched animals

Comparing the infection dynamics between nematodes inoculated immediately after hatching (0 hph) *vs* those inoculated at 6 hph (Figure 1B) revealed that the first wave of viral infection was substantially higher and prolonged in time on nematodes inoculated at 6 hph. Specifically, the maximum viral load reached (2.6 ±0.8)×10^7^ molecules RNA2/ng total RNA at ∼14 hpi for the 6 hph group, compared to (9 ±3)×10^6^ molecules after ∼12 hpi for the 0 hph group. Data in Figure 1B, fitted to a GLM (Gamma distribution, log-link function; hph as fixed factor and hpi as covariable), confirmed a significant and large-magnitude difference between the two inoculation conditions (χ^2^ = 27.127, 1 d.f., *P* < 0.001, 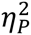 = 0.473).

This difference suggests that host physiological state at the time of infection, particularly proximity to molting, critically influences OrV replication dynamics. The enhanced viral replication in animals infected closer to a molt could indicate that the host’s capacity to restrict viral proliferation is transiently altered during this developmental transition. This observation aligns with the resource allocation principle, which states that trade-offs arise between competing functions due to limited resources^9,10^. Consequently, resources invested in producing new cuticle might not be available or be diverted from defense, resulting in impaired immune responses during molting periods.

### Late L1-inoculated nematodes exhibit a higher number of infected intestinal cells and greater luminal viral egress than early-inoculated ones

To determine whether the larger wave of viral infection observed in nematodes infected at 6 *vs* 0 hph corresponded to a greater number of infected intestinal cells, we performed smiFISH on animals collected at 12 hpi (the approximate peak of viral load for both conditions. We observed larvae with infected intestinal cells, and the presence of viral RNA in the intestinal lumen (representative images shown in Figure 2A).

**Figure 2.**
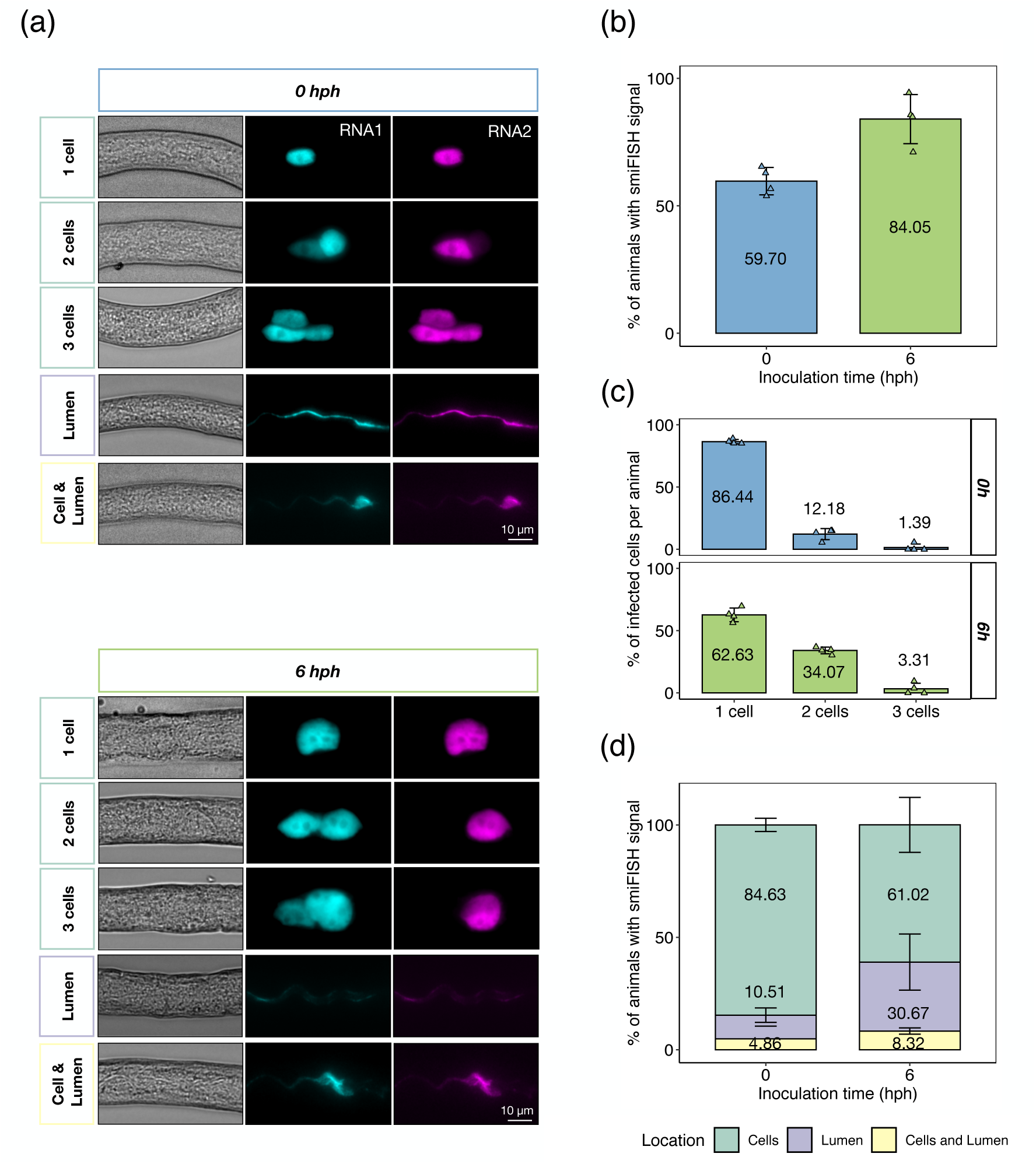
Identification of infected cells and viral presence in the intestinal lumen. **(a)** Representative smiFISH micrographs of viral RNA within cells (green boxes), on the lumen (purple box), or in both (yellow box), for both 0 and 6 hph inoculated nematodes. Signal of RNA1 is shown in cyan, and signal of RNA2 in magenta. Samples were collected at 12 hpi. **(b)** Percentage of nematodes with smiFISH signal. Nematodes inoculated at 0 hph are shown in blue, and nematodes inoculated at 6 hph are shown in green. Points represent biological replicates (each one corresponding to a plate of animals, each plate containing ∼40 animals). Mean values are shown. Error bars represent ±1 SD. **(c)** Percentage of animals according to the number of infected intestinal cells. Nematodes inoculated at 0 hph are shown in blue, and nematodes inoculated at 6 hph are shown in green. Error bars represent ±1 SD. **(d)** Percentage of smiFISH signal on cells (in green), lumen (in purple), or both (in yellow), in 0 hph and 6 hph inoculated nematodes. Error bars represent ±1 SD. On this experiment, four replicates were done for each condition, ∼40 animals per replicate.

Data (Figure 2B), fitted to a GLM (Binomial distribution, logit-link; hph as fixed factor), reveled that the frequency of nematodes with viral RNA signal was significantly higher on animals inoculated at 6 hph (0.84 ±0.03) compared to nematodes inoculated at 0 hph (0.60 ±0.05) (χ^2^ = 18.502, 1 d.f., *P* < 0.001, 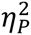 = 0.251). Furthermore, the number of intestinal infected cells per nematode was also significantly higher on 6 hph inoculated animals than in those inoculated at 0 hph (Figure 2C; Pearson homogeneity test: χ^2^ = 13.206, 2 d.f., *P* = 0.001; magnitude of the effect: Cramér’s *V* = 0.271). Specifically, 0.38 ±0.09 (LaPlace estimator ± adjusted Wald margin of error) of the 6 hph-inoculated nematodes had two or more infected intestinal cells, compared to 0.15 ±0.08 for 0 hph-inoculated nematodes.

We next assessed the spatial distribution of infection, specifically whether viral RNA localized exclusively within intestinal cells, in the lumen, or in both. The distribution of viral RNA differed significantly between animals inoculated at 0 and 6 hph (Figure 2D; χ^2^ = 15.404, 2 d.f., *P* = 0.001, *V* = 0.255). As previously reported^8^, most animals (0.84 ±0.08) inoculated at 0 hph retained the virus exclusively within intestinal cells, with only 0.11 ±0.07 of the animals showing only smiFISH signal in the lumen (all viral particles already egressed). By contrast, when inoculation took place 6 hph, the frequency of animals with only intracellular location dropped to 0.61 ±0.08. In this case, the frequency of animals with only luminal signal increased to 0.31 ±0.07, suggesting a faster completion of the intracellular infection cycle and egression to the lumen.

Molting is a highly dynamic process involving extensive physiological remodeling. Because a new cuticle must be synthesized, cellular trafficking is a fundamental aspect of this period^11–13^. This highly dynamic environment taking place during molting periods could explain the higher luminal viral signal found in animals infected near a molting stage, and the higher number of infected intestinal cells found on these animals in comparison to the ones found on freshly hatched nematodes, as the virus could potentially have more opportunities to enter and exit the intestinal cells. As OrV exits intestinal cells in a nonlytic manner^14^ (likely through exocytosis; see Figure 2A in Castiglioni et al.^8^), and considering the highly vesicular transport happening throughout the nematode’s development^15^, OrV might also sequester cellular resources, including vesicular components. Such sequestration could potentially disrupt developmental timing, adding another layer to the interplay between viral infection and host development.

### Viral infection disrupts the expression of developmental and molting-related genes, especially during the first two larval stages of *C. elegans*

As postembryonic development in *C. elegans* is highly influenced by external factors^16,17^ and can be affected by interactions with bacterial^1,2^ and fungal^3^ pathogens, we investigated the potential impact of OrV infection on *C. elegans* development. To this end, we re-analyzed transcriptional data from infected and control animals throughout larval development (NCBI SRA PRJNA1037048)^8^ and identified a set of genes exhibiting an oscillatory expression profile synchronized with molting periods. Notably, these genes (many of which belonged to molting-related functions; Table S1) follow a periodic on/off pattern that is altered in infected animals (*i.e*., compare differences between L2 - L3 phases, Figure 3A), yet converged to a similar state by the end of larval development L4 (*i.e*., at the later timepoints of the infection, Figure 3A).

**Figure 3.**
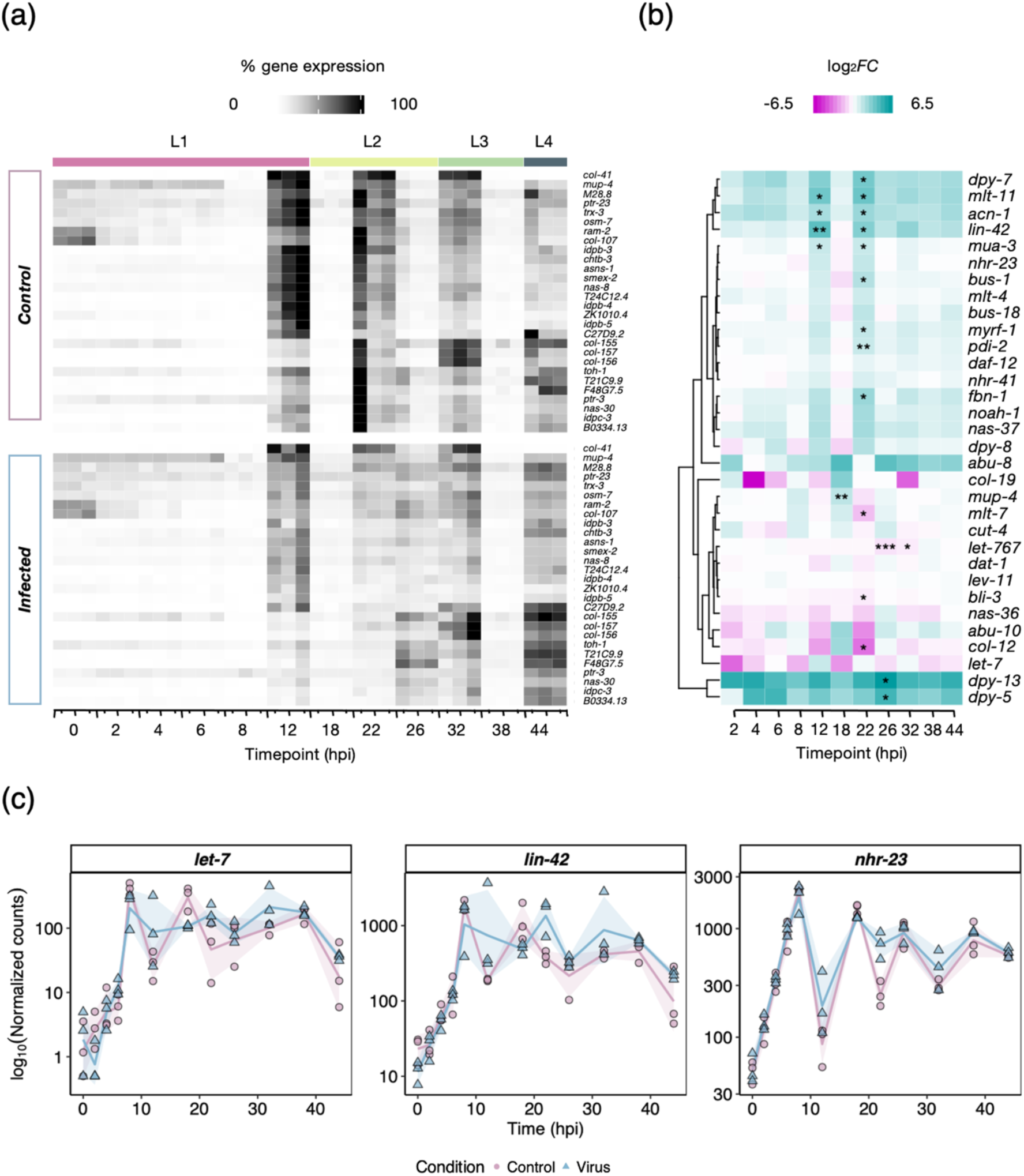
Impact of viral infection on the expression of development-related genes. **(a)** Heatmap showing the percentage of gene expression of genes with on/off cycling-like profile that matches molting times. Gradient of expression goes from white (no gene expression) to black (100% of gene expression). Recollection times (hpi) are shown on the abscissa. Three replicates (each column being one replicate) per time point were done. The developmental stages corresponding to each recollection time point are shown on the upper part of the heatmap. **(b)** Heatmap displaying the log_2_*FC* of DEGs in response to infection related with developmental and molting processes. All significant likelihood-ratio test model DEGs are shown. Additionally, significant DEGs are marked with asterisks at the specific time points in which they are significant. Recollection time points (hpi) are shown at the abscissa. Gradient of log_2_*FC* goes from magenta for downregulation to turquoise for upregulation in response to infection. (c) Expression profiles of developmental-related genes *let-7*, *lin-42* and *nhr-23* along time. Expression profiles of control animals are shown in purple circles and lines, while blue triangles and lines indicate infected animals. Shadowed areas represent ±1 SD. Three replicates per timepoint were done.

Furthermore, we found that many genes associated with development and molting were differentially expressed in response to viral infection. This includes the steroid dehydrogenase *let-767*^8^, thought to play an important role on sterol uptake^18^, an important process for extracellular matrix remodeling. Gene disruption in response to infection seemed to be especially significant during the late L1 - L2 larval stages (Figure 3B; Table S1). These findings indicate that OrV infection alters transcriptional programs linked to larval development, suggesting potential effects on molting timing.

Many *C. elegans* genes exhibit oscillatory expression patterns synchronized with the molting cycle^19–21^. Among these genes, key developmental regulators (such as the heterochronic gene *lin-42*^8,22^, the nuclear receptor *nhr-23*^23^, or the microRNA *let-7*^24^) are known to play important roles in the periodicity and control of molting timing. Mutants on these genes display disruptions in seam cell fate, causing defects on the time in which molting occurs and the number of molts happening during life, and resulting in precocious or retarded developmental phenotypes^25–27^. When analyzing the particular expression profiles of these three developmental genes on both control and infected nematodes (Figure 3C), we found that their temporal expression was significantly altered in infected animals in comparison to control ones. In the three cases the effect was due to an overall difference among both temporal expression patterns, although of small magnitude (tests of infection status by hpi interaction: χ^2^ ≥ 31.740, 11 d.f., *P* ≤ 0.001, 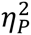 ≤ 0.019). In the case of *lin-42* and *nhr-23*, additional small yet significant effects of infection *per se* were also observed (tests of infection status: χ^2^ ≥ 6.711, 1 d.f., *P* < 0.001, 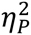 ≤ 0.003), with the average level of expression being higher in infected nematodes (grand mean 272 ±20 for *lin-42* and 496 ±18 for *nhr-23*) than in the corresponding controls (207 ±15 for *lin-42* and 402 ±15 for *nhr-23*). Altogether, these observations suggest that OrV infection indeed disrupts key aspects of nematode developmental programming.

### Virus infection shortens the L1 intermolt and advances the first molt but extends the L2 intermolt, ultimately resynchronizing developmental timing

To investigate the specific effects of OrV infection on *C. elegans* developmental timing, animals with the reporter *sevIs1[Psur-5::luc+::gfp]* (constitutively expressing luciferase) were inoculated at 0 and 6 hph. Bioluminescence assays were then performed to monitor development and molting^28^. Figure 4A shows an illustrative example of luminescence dynamics along larval development, where decreases in signal intensity correspond to molt phases. Figure 4B summarizes experimental results. The duration of the intermolt phases (Figure 4C) and molt phases (Figure 4D) were fitted to two generalized linear mixed models (GLMM) (Normal distribution, identity-link function; infection status and inoculation time as inter-subject orthogonal fix factors and phases as intra-subject random factor).

**Figure 4.**
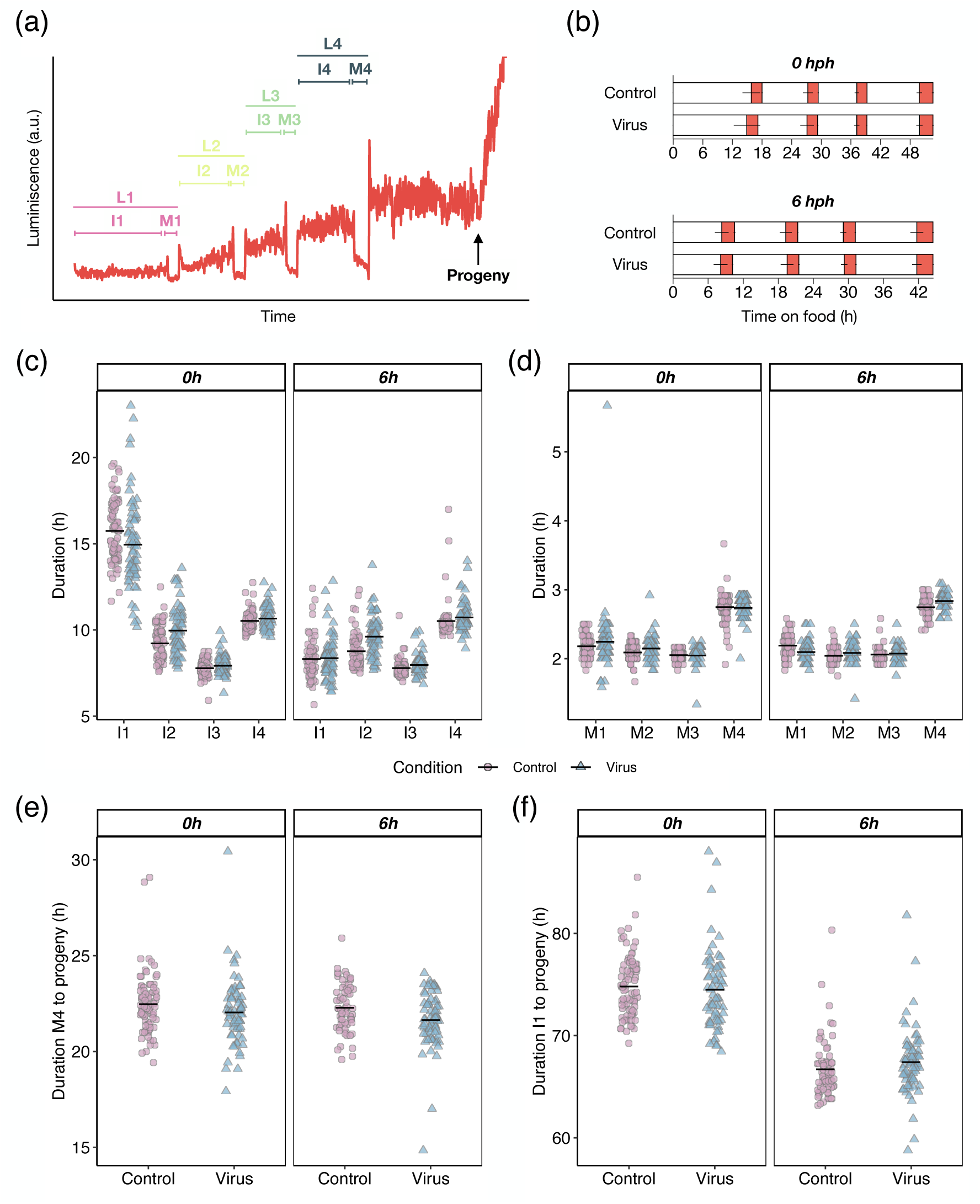
Impact of viral infection on *C. elegans* development. **(a)** Output example of the luminometry assay results. Bioluminescence signal through time (in red) of an individual nematode. Signal decays every time a molting period begins, and an exponential signal appears once offspring has hatched. Intermolt and molt stages, as well as overall larval stages (considered as both the intermolt and the molt periods) are shown. **(b)** Average duration of development for N2 animals grown at 20 °C. Red boxes represent the molts as inferred by low luciferase signal. Error bars represent the SD of the duration of each interval. **(c)** Duration of intermolt stages (I1 - I4). **(d)** Duration of molts (M1 - M4). **(e)** Time from the last molt (M4) until hatching of progeny. **(f)** Time from the first intermolt period (I1) until hatching of progeny. In panels (c) - (f) mean value are represented by black horizontal bars. Control nematodes are represented by purple circles while infected ones are represented by blue triangles.

Focusing first on intermolt phases duration (Figure 4C), infection status had a significant overall effect yet of small in magnitude in the duration of the intermolt phases (Table S2: *P* = 0.040, 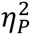 = 0.015). Not surprisingly, inoculation time has a significant effect of large magnitude (Table S2: *P* < 0.001, 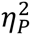 = 0.591). Remarkably, 0 hph-inoculated nematodes exhibited a shortened L1 intermolt (I1) (15 ±3 h) and initiated the first molt approximately 50 min earlier compared to control nematodes (16 ±2 h). Conversely, these nematodes displayed a lengthened L2 intermolt (10 ±1 h), extended by approximately 45 min compared to controls (9 ±1 h). Later development continued at the same pace on both infected and control nematodes.

Regarding the molt phases (Figure 4D), their duration was not affected by either infection or inoculation time (Table S3: *P* ≥ 0.079, 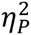 ≤ 0.011). All molts lasted approximately the same time on both infected and control nematodes. However, the final molt phase was significantly larger (2.8 ±0.2 h) than the previous three (average of 1.8±0.4 h) regardless whether animals were infected or not.

Figure 4E shows the duration from the end of the last molt (M4) to reaching sexual maturity and production of the earliest progeny. A GLM analysis (Normal distribution, identity link-function; infection status and inoculation time as orthogonal fix factors) revealed a significant, yet minor, effect of infection (χ^2^ = 8.783, 1 d.f., *P* = 0.003, 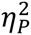 = 0.003). Infected animals reached sexual maturity ∼36 ±8 min sooner after M4 than non-infected animals (21.8 ±0.1 h *vs* 22.4 ±0.1 h).

Finally, Figure 4F illustrates the duration from the first intermolt phase and the time of sexual maturity. These data were fitted to the GLM described in the previous paragraph. As shown above, the total duration of larval development does not depend on the infection status (χ^2^ = 0.233, 1 d.f., *P* = 0.629), as the initial delay observed in infected animals is later on compensated. However, a highly significant effect, of very large magnitude, of the inoculation time has been observed (χ^2^ = 246.324, 1 d.f., *P* < 0.001, 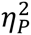 = 0.579), with animals infected at 0 hph reaching sexual maturity in 74.6 ±0.3 h, while those infected at 6 hph take 67.1 ±0.3 h.

Altogether, these results demonstrate that OrV infection transiently disrupts early larval timing but is later compensated for, resulting in synchronized overall development.

## DISCUSSION

The observation that early OrV infection accelerates the first molt, followed by a compensatory lengthening of the L2 intermolt, suggests a plastic developmental response. Prior reports indicate that *C. elegans* may trigger a sleep-like state via antimicrobial peptides acting on the same neurons that govern molting-associated quiescence, likely to bolster survival^29^. Antimicrobial effectors have also been shown to be differentially expressed in response to OrV infection^7^. Wu et al.^30^ suggested that sleep can be induced from many, perhaps all, tissues that experience energetic stress. These tissues include the intestine, the primary OrV infection site. This connection is particularly relevant as OrV infection is associated with decreased ATP levels^31^ and induces a quiescent behavior that increases survival^30,31^. The nematode’s strategy to accelerate the beginning of molting could then be an adaptive response to induce sleep-like states that enhance defense against viral infections. However, this hypothesis needs to be carefully tested. We observed that infected nematodes lengthened the L2 intermolt and resumed normal development to perfectly match the rest of developmental events relative to control non-infected nematodes. This phenomenon is reminiscent of “temporal scaling” whereby the system recalculates individual developmental events to match the overall timeline despite environmental disruptions^16^. This developmental plasticity may represent an important mechanism that allows *C. elegans* to respond to pathogenic challenges while preserving its developmental program.

## CONCLUSIONS

In this work, we have explored the interplay between OrV infection and *C. elegans* larval development, focusing primarily on (*i*) the role of molting on OrV replicative dynamics and (*ii*) the impact of OrV infection on *C. elegans* developmental timing.

Regarding the first aspect, we demonstrated that OrV maintains the ability to replicate inside intestinal cells even when the nematode is going through a molting stage, and concluded that the decay on viral load at ∼12 hpi must be explained by other factors other than entering the molting periods. Our results, in fact, suggest that molting could be impairing the nematode’s immune system functioning.

Turning to our second objective, we detected a desynchrony in developmental gene expression in infected *vs* uninfected nematodes. In freshly hatched infected nematodes, animals accelerated the start of the first molt and lengthened the L2 intermolt to resume normal development.

Overall, our results deepen our understanding of OrV virus dynamics and reveal how OrV infection can affect *C. elegans* development. Further research will be needed to dissect the molecular mechanisms underlying OrV dynamic infection profile and to clarify the biological significance of infection-induced developmental alterations in this model organism.

## RESOURCES AVAILABITLITY

### Lead contact

Further information and request for resources and reagents should be directed to and will be fulfilled by the lead contact, Santiago F. Elena.

### Materials availability

This study did not generate new unique reagents.

### Data and code availability

All data reported in this paper are accessible at Zenodo (https://10.5281/zenodo.15479384).

## ACKNOWLEDGMENTS

We thank Francisca de la Iglesia for excellent technical assistance. This work was supported by grants PID2022-136912NB-I00 funded by MCIU/AEI/10.13039/501100011033 and by “ERDF a way of making Europe”, and CIPROM/2022/59 funded by Generalitat Valenciana to S.F.E. I.M. was supported by grant PRE/2020-094661 funded by MCIU/AEI/10.13039/501100011033 and by “ESF investing in your future”. V.G.C. was supported by grant FJC2021-047264-I funded by MCIU/AEI/10.13039/501100011033 and by NextGenerationEU/PRTR. M.J.O-U. was supported by grant FPU2019/05246 funded by MCIU/AEI/10.13039/501100011033 and by “ESF investing in your future”. The work of M.O. was supported by grant PID2022-139009OB-I00 funded by MCIU/AEI/10.13039/501100011033 and by “ERDF a way of making Europe”. R.G. was supported by a Pasteur-Roux-Cantarini fellowship of Institut Pasteur.

## AUTHORS’ CONTRIBUTIONS

I.M., conceptualization, data curation, formal analysis, investigation, visualization, writing-original draft, writing-review, and editing. V.G.C., conceptualization, investigation, supervision, writing-review, and editing. M.J.O-U., conceptualization, formal analysis, investigation, writing-review, and editing. M.O. conceptualization, resources, supervision, writing-review, and editing. R.G., supervision, writing-original draft, writing-review and editing. S.F.E., conceptualization, formal analysis, funding acquisition, project administration, resources, supervision, writing-review, and editing. All authors gave final approval for publication and agreed to be held accountable for the work performed therein.

## SUPPLEMENTAL INFORMATION

**Figure S1. Schematic picture of the life cycle of *C. elegans* at 20 °C.**

**Table S1. List of differentially expressed genes related with developmental and molting relating processes.**

**Table S2. Repeated measures ANOVA summary table for the analysis of the duration of inter-molting stages.**

**Table S3. Repeated measures ANOVA summary table for the analysis of the duration of molting stages.**

**Table S4. List of RT-qPCR primers and smiFISH probes used on this work.**

## STAR METHODS

### *C. elegans* strains, maintenance and synchronization

All experiments regarding the characterization of OrV infection dynamics were done using the ERT54: *jyIs8[Ppals-5p::GFP; myo-2p::mCherry]X* strain, which is derived from the N2 Bristol and carries a fluorescent reporter for the activation of the intracellular pathogen response (IPR)^32^. Bioluminescence assays were done using the MRS387: *sevls1[Psur-5::luc+::gfp]X* strain^33^. Strains were maintained at 20 °C on nematode growth medium (NGM) plates seeded with *Escherichia coli* OP50 as food source^34,35^. For all experiments, animals were synchronized by carefully washing egg-containing plates with M9 buffer (0.22 M KH_2_PO_4_, 0.42 M Na_2_HPO_4_, 0.85 M NaCl, 0.001 M MgSO_4_) to eliminate adult nematodes and larvae and retain only eggs on the plates. After a 1 h interval, plates were washed again with M9 buffer to gather the larvae that had hatched during that time. Successive 1 h interval washing rounds were done in case more animals were needed.

### OrV stock, quantification, and inoculation procedure

SFE2: *drh-1(ok3495)IV; mjIs228[myo-2::mcherry::unc54; lys-3p::eGFP::tbb-2]?* animals in N2 background, or JU2624: *[myo-2::mcherry::unc54; lys-3p::eGFP::tbb-2]IV* animals in JU1580 background were used for viral stock preparation. Nematodes from either strain were soaked into a 500 μL aliquot containing viral stocks for 1 h. The volume was then divided among ∼30, 9 cm NGM plates and nematodes were grown at 20 °C for around one week or less, until plates ran out of food. Animals were then washed with 4.5 mL of M9 buffer each 3 or 4 plates and the volume was collected in 15 mL tubes. Nematodes were vortexed and centrifuged at 1350 rpm for 2 min. Supernatant was recovered and centrifuged twice at 14000 rpm for 5 min to eliminate any bacterial contamination. Liquid supernatant was then filtered through a 0.2 μm syringe filter and distributed in 500 μL aliquots for storage at −80 °C.

For viral stock quantification, viral RNA was extracted using the Viral RNA Isolation kit (NYZ tech, Portugal). Stock was quantified by RT-qPCR, using RNA2 as target. The primers used were oVG15_RNA2_qPCR_2_F and oVG16_RNA2_qPCR_2_R (Table S4). A standard curve method was used for quantification. Six 1/10 serial dilutions were used for the standard curve starting with a concentration of 1.25×10^9^ molecules/μL. For the pristine RNA used to make the standard curve, cDNA of OrV was obtained using Accuscript High Fidelity Reverse Transcriptase (Agilent, USA) and reverse primers at the 3’ end of the virus. Approximately 1000 bp of the 3’ end of RNA2 were amplified using forward primers containing 20 bp coding the T7 promoter and DreamTaq DNA Polymerase (Thermo Fisher, USA). The PCR products were gel purified using MSB Spin PCRapace (Invitek Diagnostics, Germany) and an *in vitro* transcription was performed using T7 Polymerase (Merck, Germany). The remaining DNA was then degraded using DNAse I (Life Technologies, USA). RNA concentration was determined by NanoDrop (Thermo Fisher, USA) and the number of molecules per μL was determined using the online tool EndMemo RNA Copy Number Calculator (https://www.endmemo.com/bio/dnacopynum.php). The primers used were oVG7_OrV_RNA2_3’_F, and oVG8_OrV_RNA2_3’_R targeting RNA2 (Table S4).

For the experiments regarding the study of viral dynamics, 6 cm NGM plates were inoculated with 60 μL of viral stock of a concentration of 4.4×10^7^ molecules/μL of OrV by pipetting the viral stock on top of the bacterial lawn containing the nematodes. For samples that were going to be collected up to 12 hours post-hatching (hph), 700 nematodes per plate were inoculated. From 12 hph until 24 hph, 500 nematodes per plate were inoculated. A minimum of three replicates were done for each timepoint.

For the bioluminescence assay, after synchronization nematodes were washed and transferred to empty plates containing M9 buffer liquid. Nematodes were pipetted one by one (on a volume of ∼2 μL) and placed on 96-well plates. One nematode was placed on each well. Ten μL of viral stock were inoculated on each well by pipetting. Four replicates were done for these experiments, each replicate being considerate as an entire 96-well plate. On each plate, a maximum of 24 nematodes were transferred for each condition (virus-inoculated nematodes at 0 hph and 6 hph, and control non-inoculated nematodes placed at 0 hph and 6 hph).

### RNA extraction and RT-qPCR

Virus-inoculated and control nematodes were washed with 1600 μL of phosphate-buffered saline (PBS)-0.05% Tween at the designated times. Nematodes were centrifuged at 1350 rpm for 2 min and the supernatant was discarded. Two additional rounds of washing were performed for each sample using ∼620 μL of PBS-0.05% Tween. Samples were then quick frozen in liquid N_2_ and stored at −80 °C until extraction procedure was initiated.

Five hundred μL of Trizol (Invitrogen, USA) were added to the nematode pellet containing a volume of ∼100 μL of PBS-0.05% Tween. Samples were then vortexed, and five cycles of freeze-thawing were performed (from liquid N_2_ to 37 °C). Samples could be frozen again after this step until extraction protocol was continued. Five rounds of 30 s of vortex followed by 30 s of rest were then performed. One hundred μL of chloroform were added to the aliquots, tubes were shaken for 15 s, and then let rest for 2 min at room temperature. Samples were centrifuged for 15 min at 12,000 g at 4 °C. The top layer containing the RNA was recovered and mixed with the same volume of 100% ethanol. Samples were then loaded into RNA Clean & Concentrator columns (Zymo Research, USA) and the rest of the protocol was followed according to manufacturer instructions.

RT-qPCRs were performed using Power SYBR Green PCR Master Mix (Applied Biosystems, USA) on an ABI StepOne Plus Real-time PCR System (Applied Biosystems, USA). Ten ng of total RNA per sample were loaded, and a standard curve method was used for OrV quantifications (as described for viral stock quantification). The primers used were the same as the ones used for viral stock quantification (Table S4).

### smiFISH

smiFISH protocol was adapted from previous studies^36,37^. Briefly, virus-inoculated, and control nematodes were washed with 1600 μL of PBS-0.05% Tween and centrifuged for 2 min at 1350 rpm, and supernatant was discarded. Three additional rounds of washing were performed for each sample using ∼620 μL of PBS-0.05% Tween. Eight hundred μL of Bouins fixation mix (400 μL Bouins Fix, 400 μL methanol, 10 μL β-mercaptoethanol) were then added to the samples, which were later incubated at room temperature for 30 min in a rotatory shaker. Samples were then quick frozen in liquid nitrogen and kept at −80 °C until continuation of the protocol.

The day prior to the microscopy assay, samples were gently shaken at 4 °C in a rotatory shaker for 30 min and washed 4 times with borate Triton solution (20 mM H_3_BO_3_, 10 mM NaOH, 0.5% Triton), and 5 times with borate Triton β-mercaptoethanol solution (20 mM H_3_BO_3_, 10 mM NaOH, 0.5% Triton, 2% β-mercaptoethanol), leaving the borate Triton β-mercaptoethanol incubate for at least 1 h between each wash in the last three washes. Samples were then incubated for 5 min with wash buffer A (1 mL Stellaris RNA FISH Wash Buffer A, 1 mL deionized formamide and 3 mL H_2_O). Forty probes against OrV were designed using Oligostan^36^; 22 against RNA1, and 18 against RNA2 (Table S4). The probes were annealed as follows: 2 μL of 0.83 μM probe set, 1 μL of 50 μM fluorescent labeled amplification product (FLAP) label, 1 μL NEB3, and 6 μL of diethylpyrocarbonate-treated H_2_O were incubated at 85 °C for 3 min, 65 °C for 3 min, and 25 °C for 5 min. The FLAP labels contained CAL Fluor 610 or Quasar 670 modifications at both the 5’ and 3’ ends of the sequence 5’-AATGCATGTCGACGAGGTCCGAGTGT-3’ (Biosearch Technologies, UK). One μL of annealed probe solution targeting RNA1 and 1 μL of annealed probe solution targeting RNA2 were then mixed with 98 μL of hybridization buffer (100 μL Stellaris smiFISH hybridization buffer, 25 μL deionized formamide). A volume of 100 μL of hybridization buffer containing the smiFISH probes were then added to the samples and the samples were incubated overnight at 37 °C in a rotary shaker. The following day, samples were washed with Wash Buffer A2 (1 mL of Stellaris RNA FISH Wash Buffer A, and 4 mL of H_2_O), incubated for 30 min in Wash Buffer A2 at 37°C, and then incubated for 30 min in Wash Buffer A2 containing 25 ng of 4’,6-diamidino-2-phenylindole (DAPI) at 37 °C. Samples were then incubated for 5 min in Stellaris RNA FISH Wash Buffer B, centrifuged, and resuspended in a small volume of Stellaris RNA FISH Wash Buffer B. A quantity of 0.1 ng of DAPI was added to each sample, and samples were mounted on slides using N-propyl gallate mounting medium.

Samples were imaged using Leica DMi8 microscope with Leica DFC9000 GTC scientific complementary metal-oxide semiconductor camera and objective HC PL APO 40×/0.95 CORR PH2. Images were analyzed and processed using ImageJ (Fiji)^38^. smiFISH viral fluorescent signal was used to determine the percentage of infected nematodes, the number of infected intestinal cells on each nematode, and the spatial localization of the viral fluorescent signal. Samples were collected at 12 hpi after inoculating nematodes at 0 and 6 hph. Five replicates were done for each condition, 40 animals per replicate observed.

### Bioluminescence assay and luminometry data analysis

*C. elegans* strain MRS387 constitutively and ubiquitously expresses the enzyme luciferase. Larvae form a plug of extracellular material in the buccal cavity during molting periods, ceasing pharyngeal pumping^39^. This allows for a detailed quantification of molting time using the luciferase assay, since non-feeding individuals do not ingest luciferin, resulting in the cessation of light production and a decrease in the luminometry signal. Single synchronized L1 animals were transferred into a well of a white, flat bottom, 96-well plate by manual picking as previously described. Each well contained 160 μL of NGM mixed with 5 μL of *E. coli* OP50, and 10 μL of 1 mM D-Luciferin (plus 10 μL of OrV stock for the infected nematodes). Plates were sealed with a Breathe Easier gas-permeable cover (Diversified Biotech, USA) prior to their introduction on the microplate luminometer. Luminescence was read for 1 s, at 5 min intervals, for a period of ∼100 hours with a Berthold Centro LB960 XS3 (Berthold Technologies, Germany). Experiments were performed at 20 °C inside a Panasonic MIR-154 temperature-controlled incubators (PHC Europe B.V., The Netherlands). Temperature was monitored using HOBO Data Loggers (Onset, USA).

The raw data from the luminometer was analyzed to detect the beginning and end of the molts as follows. The raw data was trend-corrected dividing by a centered moving average. The moving average was calculated for the duration of 10 h (approximated duration of each at 20 °C). To determine the timing of the molts, the trend-corrected data was converted to binary, using 75% of the value of the moving average as a threshold. The data was then evaluated for onset and offset of molting by detecting the transitions in the binarized data. Transitions from 1 to 0 corresponded to the onset of a molt, and transitions from 0 to 1 corresponded to the offset of a molt. To prevent the noise in the binarized data product of the variability introduced by the luminometry signal, the following rule was applied: onsets must be followed by “0” for at least 1 h, and offsets must be preceded by “0” for at least 1 h. The duration of each intermolt and molt stage, the time from the end of the last molt until the start in the production of progeny (characterized by an exponential increase in the luminometry signal), as well as the duration of development from the first intermolt until the production of progeny, were then calculated. Plots were done using ggplot2^40^ in R version 4.2.0^41^ under RStudio version 2022.7.1.554.

### Identification of genes with a switch-like expression pattern

DEGs previously identified (raw data from NCBI SRA PRJNA1037048) as genes with a differential gene expression profile (likelihood ratio test) between OrV infected and noninfected were used here^8^. The gene expression counts of the uninfected animals were normalized to the maximum value per gene. Hierarchical clustering of transcriptional profiles along larval development was then applied using Euclidean distance. Visual exploration of all the clusters allowed us to identify one with a clear oscillatory expression pattern.

### Statistical analysis

Generalized linear models (GLM) were fitted to the different datasets by maximum likelihood using SPSS version 30.0.0.0 (IBM, Armonk, NY). Specific model designs are introduced in the text as necessary. Bonferroni *post hoc* tests were used to pairwise comparisons among factor values. The magnitude of the effects was evaluated using the 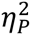 statistic (proportion of the total variability in the variable attributable to each factor in the model; conventionally, values of 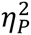≥ 0.15 are considered as large effects). Otherwise indicated, mean ±1 standard deviation will be reported. Additional statistical tests will be introduced as needed.

